# Metformin-mediated mitochondrial protection post-cardiac arrest improves EEG activity and confers neuroprotection and survival benefit

**DOI:** 10.1101/2021.10.07.460078

**Authors:** Muhammad Shoaib, Rishabh C. Choudhary, Rupesh K. Chillale, Nancy Kim, Santiago J. Miyara, Shabirul Haque, Tai Yin, Maya Frankfurt, Ernesto P. Molmenti, Stavros Zanos, Junhwan Kim, Lance B. Becker

## Abstract

Cardiac arrest (CA) produces global ischemia/reperfusion injury resulting in substantial multiorgan damage. There are limited efficacious therapies to save lives despite CA being such a lethal disease process. Surviving patients suffer extensive brain damage with mitochondrial dysfunction implicated as a major source of injury. Metformin, a first-line treatment for diabetes, has also shown promising results in other diseases and is known to interact with mitochondria. In the present study, we evaluated the therapeutic benefits of metformin administration immediately after resuscitation using a 10 min asphxyial-CA rat model. This is the first study to show that metformin treatment post-CA a) improves survival and neurologic function, b) potentiates early normalization of brain electrophysiologic activity, and c) protects mitochondrial function with a reduction in apoptotic brain injury. Overall, as an effective and safe drug, metformin has the potential to be an easily translatable intervention for improving survival and preventing brain damage after CA.

**Summary:** Brain damage post-cardiac arrest is a major cause of death; metformin protects brain mitochondria and improved survival and brain function.

## Introduction

Cardiac arrest (CA) is a major health crisis with a very high mortality rate, resulting in greater than 300,000 deaths annually in the US with the majority of patients not surviving to hospital discharge ^1^. Studies have shown that survival after CA may be dependent on the degree of brain protection after resuscitation, which has been a major limitation faced by most clinicians ^2^. With recent technological advances in advanced life support methodologies, such as extra-corporeal membrane oxygenation (ECMO) and hypothermia, there is some optimism for improving brain protection. However, these non-specific technologies may not directly confer neuroprotection, necessitating a requirement for more targeted pharmacological interventions ^3^. The brain, as well as many other organs, undergoes a variety of pathological changes post-CA, such as increased oxidative stress, altered metabolism and energy production, and dysregulated cerebral autoregulation ^4^. One major source of pathology, which may also serve as a potential target post-CA, is the mitochondria. Mitochondrial injury from ischemia/reperfusion injury (IRI) results in energy and metabolic dysfunction, along with propagation of cellular death pathways ^5^. As such, many studies have attempted mitochondrial protection post-CA using a variety of approaches, such as protecting mitochondrial membranes and lipids using SS-31 ^6^, modulating mitochondrial reactive oxygen species production (ROS) using MitoSNO ^7^, or mitigating the destabilization of mitochondrial gradients via the mitochondrial permeability transition pore complex (MPTP) using cyclosporine A ^8^. However, the mechanisms of these newer interventions, especially in the context of mitochondria, are not fully understood, posing a challenge for researchers when translating these therapies to human CA patients.

One way to circumvent some of these challenges is to repurpose a commonly used agent, such as metformin, that has a relatively safe profile in humans and is capable of targeting multiple sites which can be exploited based on the disease. The use of metformin as a first-line treatment for type II diabetes mellitus is universally adopted despite the lack of clarity in the exact mechanism(s) that confer therapeutic benefit ^9,10^. Furthermore, along with its role in diabetes, metformin has also been implicated in having anti-aging and anti-tumor properties as well as providing protection against the cardiovascular and neurological sequalae of diabetes ^10^. Furthermore, metformin is able to exert its benefits by modulating mitochondria. As such, implementation of metformin therapy after return of spontaneous circulation (ROSC) may be able to provide favorable outcomes post-CA as it may be able to directly influence one major source of injury after CA.

The therapeutic efficacy of metformin and possible mechanism of action as a treatment for cardiac arrest (CA) is not known. One study using a rat model of 9 min asphyxia-CA evaluated the effects of 2 weeks pretreatment with metformin on survival and neurological outcomes and found increased 7-day survival and reduced neurological deficit ^11^. The major therapeutic mechanism observed to confer protective outcomes was metformin-induced activation of AMPK-dependent autophagy that aided in protecting the brain from IRI. Long-term pretreatment with metformin seemed to result in more substantial AMPK activation. This suggests that chronic metformin treatment may have the potential to improve survival post-CA. Some studies on the use of metformin as a potential therapy for IRI used models of localized injury, such as stroke or myocardial infarction. Providing metformin either before myocardial infarction or before reperfusion in a murine model conferred substantial cardioprotection that was mediated by increased activation of AMPK leading to increased activation of eNOS ^12^. However, chronic rather than acute pretreatment with metformin in a murine model of middle cerebral artery occlusion conferred neuroprotection mediated by nNOS, but without increasing AMPK ^13^. This is in contrast to studies in stroke that suggest that metformin improve outcomes by promoting AMPK ^14^. However, another major mechanism of metformin is possible. Cancer and metabolic disease studies suggest that the effects of metformin are mediated by mitochondria ^15,16^. A recent meta-analysis suggested that metformin can reduce all-cause mortality in patients with coronary artery disease as compared to another antidiabetic medication ^17^. A pilot study suggests that human patients on metformin showed improved cardiac and renal function parameters after a sudden cardiac arrest (SCA) as opposed to patients not on metformin, which was confirmed in a SCA mouse model ^18^. The protective nature of chronic, pretreatment with metformin and favorable health outcomes has growing evidence. Although the complexity of the timing, dose, and mechanism of metformin’s therapeutic benefits after IRI is unclear, it is a treatment option worth exploring. In our study, we assessed the therapeutic benefits of administering metformin after CA in a rat model of 10 min asphyxial-CA by evaluating survival, neuroprotection, and potential mechanism(s) of action.

## Materials and Methods

### Animal experimental procedure for 10 min of asphyxial-cardiac arrest

All experiments were carried out in accordance with the approval of the Institutional Animal Care and Use Committee (IACUC) guidelines at Feinstein Institute for Medical Research (2016-009). Male Sprague-Dawley rats (400-500 g; Charles River Laboratory, Wilmington, MA) were housed on a 12-h light/dark cycle with free access to water and food prior to experiment. The procedures for inducing asphyxia have been previously published ^19^. In brief, rats were anesthetized with 4% isoflurane (Isothesia, Butler-Schein AHS), intubated with a 14-gauge plastic catheter (Surflo, Terumo Medical Corporation, NJ, USA), and put onto mechanical ventilation under 2% isoflurane. The left femoral artery and vein were cannulated with polyethylene catheters (PE50, BD Intramedic, USA) to measure arterial pressure and drug infusion, respectively. After surgical preparation and heparin injection (300 IU) animals were observed for the mean arterial pressure (MAP) had normalized. Baseline recording of mean arterial pressure, heart rate, end-tidal CO_2_ (ETCO_2_) were done, which continued during CA as well as 2 h post-ROSC using PowerLab and LabChart (ADInstruments, USA). The procedure for CA began with injecting vecuronium bromide (2 mg/Kg by body weight; Hospira, USA) slowly administered through the left femoral vein over 4 min interval. After 3 min of vecuronium injection, asphyxial-CA was induced by switching off the ventilator and subsequently discontinuing isoflurane. Mean arterial pressure below 20 mmHg was defined as CA ^19^. After 10 min of untreated asphyxia-CA, resuscitation was started with the resumption of ventilation under 100% oxygen, initiation of chest compressions, and a 20 μg/Kg bolus injection of epinephrine (International Medication System, Limited, USA) to achieve ROSC defined as MAP greater than 60 mmHg. Rats were then monitored on ventilation for 2 h post-ROSC. Esophageal temperature was maintained at 37 ± 0.5 by providing external heat. At 2 h post-ROSC rats were weaned from the ventilator, all the catheters were removed, and wounds were sutured. Rats were then returned to the animal housing facility and provided daily care according to the approved protocol. Animals were not included in the study if they did not achieve ROSC within 5 min after initiation of CPR. Animals were monitored for 72 h survival. Animals were then euthanized, and the whole brain was harvested for histological and biochemical analyses. For comparison with non-CA animals, sham animals were deeply anesthetized and euthanized for tissue collection according to the protocols for the respective experiments. A schematic of the experimental protocol is shown in Fig. 1.

**Fig. 1:**
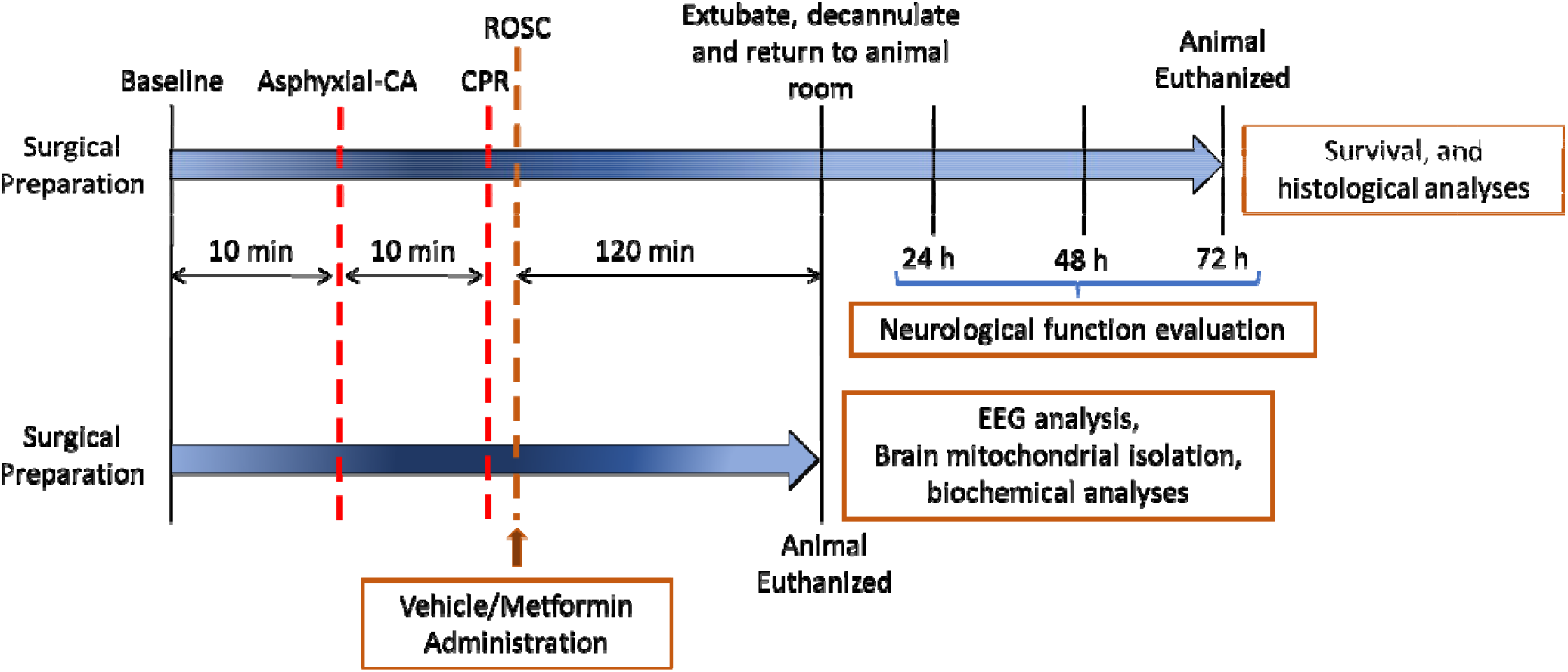
Experimental Design. Diagrammatic representations of the experimental design for rodent cardiac arrest for survival (top) and for EEG, mitochondrial isolation, and biochemical analyses (bottom).

### Experimental design for metformin administration

Rats were randomly assigned to vehicle- or metformin-treated groups. Animals received either a vehicle (saline) or metformin (EMD Millipore Corp. USA; 100 mg/kg ^20^ body weight in 2 mL saline) injection through the femoral vein just after achieving ROSC over 15-20 min using an infusion pump (Kent Scientific Corporation, USA). Blood was collected from femoral artery at baseline, ROSC, 20 min, and 40 min post-ROSC. Blood samples were then centrifuged at 1000 x g for 10 min to separate the plasma, and the collected plasma was immediately frozen and stored in -80 ° C for further biochemical analysis. For arterial blood chemistry analysis, pH, pCO_2_, pO_2_, HCO_3_, lactate, and oxygen saturation were measured at baseline, 20 min, and 40 min post-ROSC. Glucose measurement was done at baseline, ROSC, 20 min, and 40 min post-ROSC.

### Recording of Neurological Deficit Scores at 24, 48, and 72 h post-resuscitation

Rats undergone through 10 min CA and CPR after achieving ROSC were monitored for 72 h survival to record survival rate (n=20 each). To measure neurological recovery a recording of modified neurological deficit Scores (mNDS) was performed in a blinded manner using a modified version of a previously established method ^21^ (table S1). The mNDS values were monitored at 24, 48, and 72 hrs post-ROSC (mNDS, 0-500; 0, brain dead; 500, normal).

### Quantitative Electroencephalographic analysis after 10 min CA and resuscitation

We measured EEG during baseline and up to 2 h after ROSC. Briefly, before intubation and cannulation, animals were anesthetized with isoflurane and then put on a stereotaxic apparatus (Stoelting, USA). EEG electrodes implantation was performed bilaterally into the frontal and parietal cortex. Each animal had five screw electrodes (Plastics One, Roanoke, VA) cortically implanted at 2 mm lateral and 2 mm anterior or posterior to the bregma using a rat brain atlas ^22^. In addition, a ground electrode was placed 2 mm right lateral and 9 mm anterior to the bregma in the midline over the parasagittal right frontal lobe. EEG was recorded using an Intan RHS Stim/Recording 16 channel instrument (Intan Technologies, USA) during the baseline and after ROSC. Raw EEG signals were used to determine the EEG electrical activity.

The digitized electroencephalographic (EEG) signals were analyzed using MATLAB, version *9*.*0*.*0341360* (R2016a) (Natick, Massachusetts: The MathWorks Inc, 2016). After completion of animal recordings (n=6 each for vehicle- and metformin-treated groups), all EEG signals at baseline and during the post-resuscitation periods were subject to manual inspection to detect any abnormalities in the recordings. Animals were excluded if they expired shortly after achieving ROSC, and/or showed a severely abnormal EEG recording (i.e: signal/noise ratio indiscernible). After the manual inspection, all EEG signals were marked as isoelectric/suppression or burst suppression by an independent, blinded investigator ^23-25^. The criteria used for the classification is as follows: 1) Isoelectric/suppression is identified when there is no significant EEG for at least a 30 sec time-period, and 2) the criteria for EEG amplitude > 10μV to detect the bursts and burst suppression is characterized by at least 0.5 sec of no EEG activity (or EEG amplitude < 10μV) between the bursts.

After this classification, 1 min artifact-free EEG segments were chosen for baseline and at each 10 min mark during the post-ROSC period. The EEG segments were processed to remove any offsets and high frequency noise. We used a band pass filter in the range of 0.3-100 Hz. The EEG segments were subjected to burst suppression calculation. The burst suppression ratio (BSR) can be defined as the duration of the burst signal to the total duration of the signal. We computed the BSR for each channel and in each condition. In each condition, the BSR for all epochs was averaged across all channels and used in subsequent statistical analysis. The complete experimental design is shown in fig. S1.

### Histology and staining

At 72 h post-ROSC, surviving animals were placed under anesthesia and transcardially perfused using cold phosphate buffer saline (PBS, 1X, pH= 7.4). Sham animals were non-CA rats that were deeply anesthetized and euthanized after reperfusion with PBS. Whole brains from sham or experimental groups were harvested and fixed in 4% paraformaldehyde (PFA) at 4°C. Following cryopreservation in 30% sucrose solution in PBS, serial coronal sections (14 μm) containing the hippocampus were cut on a cryostat (CM1900, Leica) and collected on subbed glass slides. Nissl staining (Cresyl Violet, Acros Organics, USA), NeuN and GFAP staining (Thermofisher Scientific, USA) were performed using a previously established method ^26,27^, while TUNEL staining (Abcam, UK) was performed according to the manufacturer’s instructions. NeuN and GFAP immunostaining were performed after antigen retrieval using antigen retrieval solution (Dako, Target Retrieval Solution, pH 6, 1X, Denmark) and incubated in a pressure cooker for 5 min. After antigen retrieval, slides were incubated with 0.25% Triton X for 20 min (in 1X TBST) and afterwards blocked with 2% normal donkey serum (NDS; VWR, USA) in 1X, TBST at room temperature. After incubation, sections were incubated with mouse anti-rat NeuN (1:500) and rabbit anti-rat GFAP (1:500) antibodies prepared in 2% NDS in 1X TBST. Slides were incubated overnight at 4°C with slight agitation. For double immunostaining, slides were washed 3 times with 1X TBS and incubated with secondary antibody for NeuN (Green, AlexaFluor 488, donkey anti-mouse, 1:400) and for astrocytes (Red, AlexaFluor 594, donkey anti-rabbit, 1: 400) for 1 h at room temperature. Following incubation, slides were washed three times with 1X TBS followed by water and mounted using Fluoroshield mounting medium with DAPI (Sigma, USA).

Pyramidal neurons from the CA 1 region of the hippocampus in 4 serial coronal sections for sham, vehicle, and metformin group (n=4 each) were imaged at 40X magnification bilaterally using the BX-X800 bright field microscope (Keyence, USA). BX-X800 Analyzer (Keyence, USA) was used to semi-quantify the number of ischemic (Nissl staining) and apoptotic cells (TUNEL staining) from CA 1 hippocampal region averaged from both hemispheres. For NeuN and GFAP, hippocampal CA 1 regions were imaged using the BX-X800 fluorescent microscope (Keyence, USA). ImageJ was used to semi-quantify the neurons and astrocytes from both CA 1 hippocampal areas which were then averaged. All counting was done per focal area.

### Assessment of AMPK mRNA and protein content in brain tissue

Brain tissue was prepared for immunoblotting, as previously reported ^28^. Western blot analysis of p-AMPK α, p-AMPK β1, Total AMPK α, and Total AMPK β1/2, was conducted using brain homogenate. Sham animals were non-CA rats that were deeply anesthetized and euthanized. Briefly, sham (n=3), vehicle- and metformin-treated rats (n=4 each) were transcardially perfused using cold phosphate buffer saline (PBS, 1X, pH= 7.4) and whole brain was collected. Brains were flash frozen and pulverized into powder. Pulverized brain (20 mg) was taken and 100 μL of RIPA buffer (Sigma, USA) with Pierce Protease Inhibitor (ThermoFisher Scientific, USA) and 1 mM sodium orthovanadate (Sigma, USA) was added; the brain was sonicated 5 times for 10 sec in ice. Protein concentration from rat plasma was quantified using bicinchoninic acid protein assay (ThermoFisher Scientific, USA). Samples were diluted with Laemmli SDS Sample Buffer and heated at 95-100°C for 5 min followed by centrifugation at 16,000 x g at 4°C for 2 min. Samples were resolved using 4–20% precast polyacrylamide gel electrophoresis and transferred to polyvinylidene fluoride membrane (Bio-Rad Laboratories, Hercules, CA). After blocking, the membrane was incubated overnight at 4°C with the primary antibodies for p-AMPK α, p-AMPK β1, Total AMPK α, and Total AMPK β1/2 (1:1000; Cell Signaling Technology, Beverly, MA). Detection used anti-rabbit horseradish peroxidase–conjugated secondary antibody (1:2000; Cell Signaling Technology, Beverly, MA). Membranes were detected using enhanced chemiluminescence detection method (ThermoFisher Scientific, USA) and imaged using ChemiDoc MP Imaging System and quantified using Image Lab Software (Bio-Rad Laboratories, Hercules, CA). Protein content was quantified as the ratio of p-AMPK/total AMPK for the subunit evaluated.

The expression of AMPKα1, AMPKα2, and AMPKβ mRNA was measured by RT-qPCR employing TaqMan probes using the CFX-96 real-time PCR system (Bio-Rad, Hercules, CA, USA). The RT-qPCR TaqMan assay was carried out with the BrightGen HER2 RT-qDx assay kit (Syantra, Calgary, Canada) according to the manufacturer’s recommendations. Real-time PCR amplification for mRNA was performed using a total volume of 20 μL (10 μL of 2x Thunderbird probe qPCR mix (Toyobo, Osaka, Japan), 5 μL of primer and TaqMan probe mixture, 2 μL of template cDNA, and 3.0 µl nuclease free water). PCR cycle conditions were 95°C for 3 min followed by 15 sec at 95°C and 30 sec at 55°C for 40 cycles. To avoid false negatives due to degradation of mRNA, glyceraldehyde-3-phosphate dehydrogenase (GAPDH), was used as an endogenous control. Target mRNA expression (fold change) relative to GAPDH was automatically calculated using the comparative Ct method by CFX Manager Software v1.6 (Bio-Rad Laboratories, Hercules, CA) or Genex Software (Bio-Rad Laboratories, Hercules, CA) and the cut-off value for distinguishing between positive and negative results was a relative HER2 mRNA expression level of 100.

### Isolating brain mitochondria

Brain mitochondria were isolated from sham (non-CA) animals (n=6) after they were deeply anesthetized and euthanized as well as from vehicle- and metformin-treated rats (n=6 each) after 2 h post-ROSC using a modified differential centrifugation method as published previously ^29^. All the procedures were performed at 4°C. Briefly, brain was placed into mitochondrial isolation buffer (MESH) composed of 210 mM mannitol, 70 mM sucrose, 10 mM Hepes, 0.2 mM EGTA, at a pH of 7.3. Brains with the spinal cord removed were blotted dry, weighed, and placed into MESH buffer with 0.2% w/v fatty acid free-bovine serum albumin (MESH-BSA). Brain was minced well and 10 mL/g tissue of MESH-BSA was added. The tissue was subsequently homogenized using teflon/glass motor-driven homogenizer (Model BDC2010, Caframo Lab Solutions, Ontario, Canada) with 8 strokes. The homogenate was centrifuged at 5600 x g for 1 min; the supernatant was collected in a polycarbonate tube, and the pellet was dissolved in MESH-BSA and homogenized again. The supernatant was pooled and centrifuged at 11000 x g for 6 min. The remaining loose pellet was suspended with 20 mL of 12.5% percoll in MESH (v:v) and centrifuged at 11000 x g for 6 min. The supernatant was gently decanted with pipets without disturbing the mitochondria pellet, which was suspended in 0.2 mL MESH per g of tissue. Mitochondria concentrations were determined by the BCA assay and expressed as mg mitochondrial protein/g tissue.

### Biochemical analysis of complex I, IV activity, cytochrome c, and protein carbonyl in whole brain and brain isolated mitochondria

Sham, vehicle- and metformin-treated rats were transcardially perfused using cold phosphate buffer saline (PBS, 1X, pH= 7.4) and whole brain was collected. Brains were flash frozen and pulverized into powder, which was used to determine protein carbonyl concentration (Cayman Chemical, USA) and complex 1 activity (Abcam, Cambridge, UK) as per manufacturer’s protocol. Complex I, IV, and cytochrome c (Abcam, Cambridge, UK) levels were measured using isolated brain mitochondria as per manufacturer’s protocol.

### Statistical Analyses

Data for continuous variables are presented as mean ± standard error of the mean (SEM). Categorical data are presented as frequencies with proportions. For hemodynamic parameter measurements and arterial blood chemistry analysis, repeated measures two-way analysis of variance (ANOVA) followed by either Tukey’s or Sidak’s correction for post-hoc comparison to compare differences in the same group, or between groups, respectively. The proportion of rats surviving until 72 hours was evaluated using the Gehan-Breslow-Wilcoxon test to compare the survival curves between the two groups. For histologic comparison of brains isolated from sham and the surviving animals from the vehicle- and metformin-treated groups, one-way ANOVA followed by Tukey’s multiple comparisons test was used to compare differences among groups. For EEG analysis, the BSR was calculated using Repeated Measured two-way ANOVA with Bonferroni’s multiple comparisons test, and the time to reach 50% baseline BSR was calculated using unpaired Student’s T-test. For other analyses, an unpaired two-tailed Student’s t-test, Mann-Whitney U test, or one-way/two-way ANOVA was used for continuous variable comparison as appropriate. Statistical significance was set at p<0.05. GraphPad Prism 9.1 (GraphPad Software Inc., La Jolla, CA, USA) was used for statistical analyses.

## Results

### Metformin administration improves survival outcomes and mitigates neurological dysfunction after 10 min CA and resuscitation without changes in hemodynamic or blood chemistry parameters

No significant differences were observed in baseline characteristics between vehicle- and metformin-treated rats in body weight (469.70 ± 7.67 vs 468.70 ± 7.11 g; P=0.85); time to achieve CA (203.10 ± 5.42 vs. 192.40 ± 6.27 s; P=0.26); and time to achieve ROSC (63.75 ± 5.07 vs. 69.95 ± 4.96 s; P=0.29). Table S2 summarizes the general physiological characteristics of rats used in the experiment. A singular administration of metformin immediately after ROSC demonstrates a significantly improved survival benefit at 72 h as compared to vehicle-treated rats (65.0% v 45.0%; P=0.02; Fig. 2). Neurological evaluation conducted at 24, 48, and 72 h after ROSC demonstrates that, of the surviving rats, metformin-treated rats have significantly improved neurologic function as compared to vehicle-treated rats (P<0.01 at 24 h; P< 0.05 at 48 h; and P<0.01 at 72 h).

**Fig. 2:**
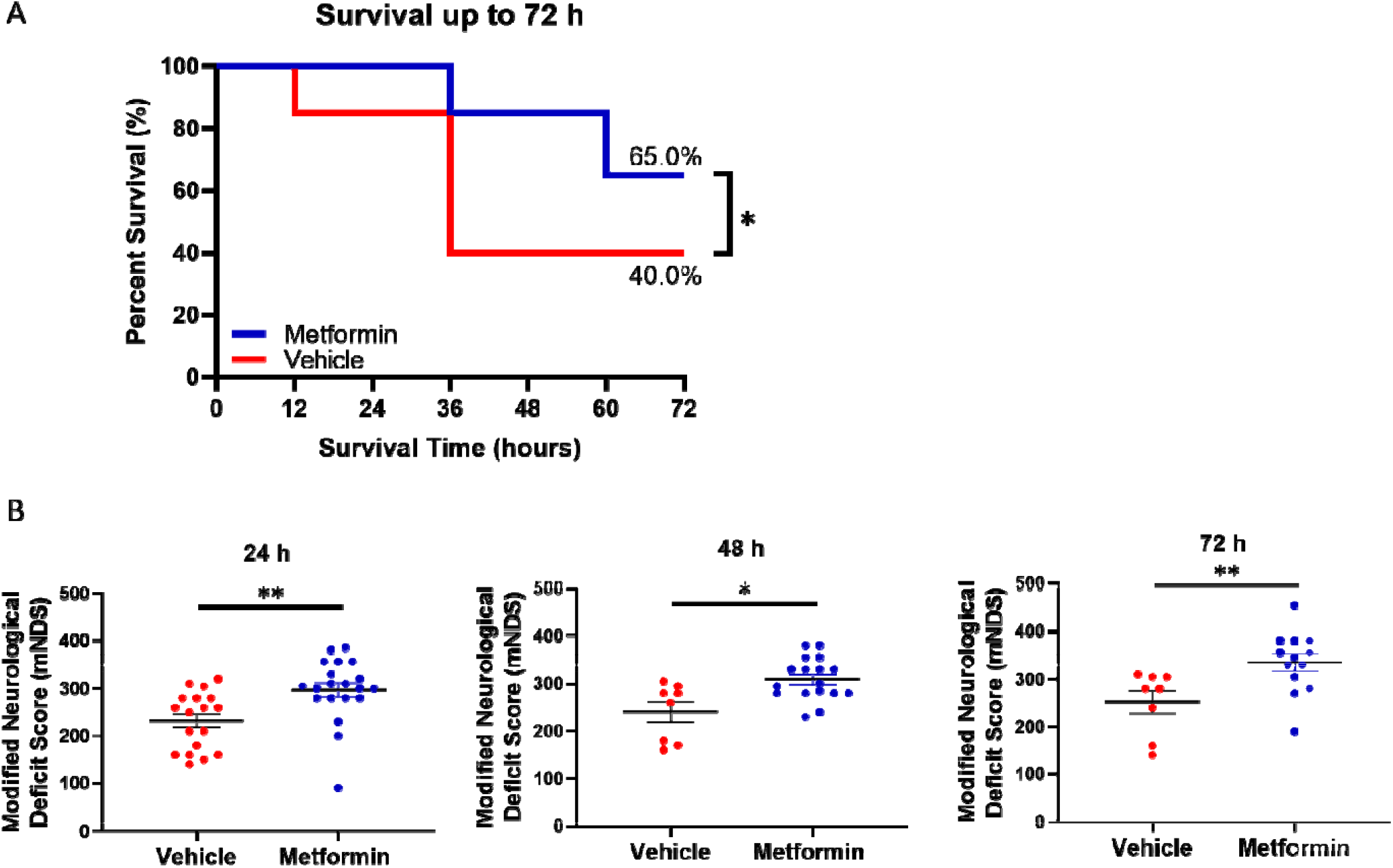
Survival and neurological function. Analysis of survival (A) and neurological function (B) up to 72 h post-ROSC between vehicle- and metformin-treated groups after 10 min cardiac arrest and resuscitation. *P<0.05 and **P<0.01.

Evaluation of plasma glucose and lactate concentrations post-ROSC in both vehicle- and metformin-treated groups shows that no significant difference between groups was observed in blood glucose and lactate levels post-ROSC as compared to their respective baselines (Fig. 3A). No significant changes in hemodynamic parameters were observed between groups except in esophageal temperature at 60 and 90 in post-ROSC (Fig. 3B). Although the vehicle group displayed temperatures at the upper limit of normal, they were still within the parameters of the experimental design. Intragroup hemodynamic changes were unremarkable as they followed a similar trend as expected for CA over time (table S3). Table 1 shows the intragroup arterial blood chemistry analysis within the vehicle- and metformin-treated groups. Although there were substantial changes over time post-ROSC in both groups, most parameters showed a similar degree of alteration in both groups. No major changes in hemodynamic parameters and blood chemistry as a function of metformin treatment reinforce the safety of this intervention for CA. Taken together, our data demonstrates that metformin administration after ROSC seems to improve survival and protect brain function without significant alterations in blood chemistry as compared with vehicle treatment.

**Table 1.**
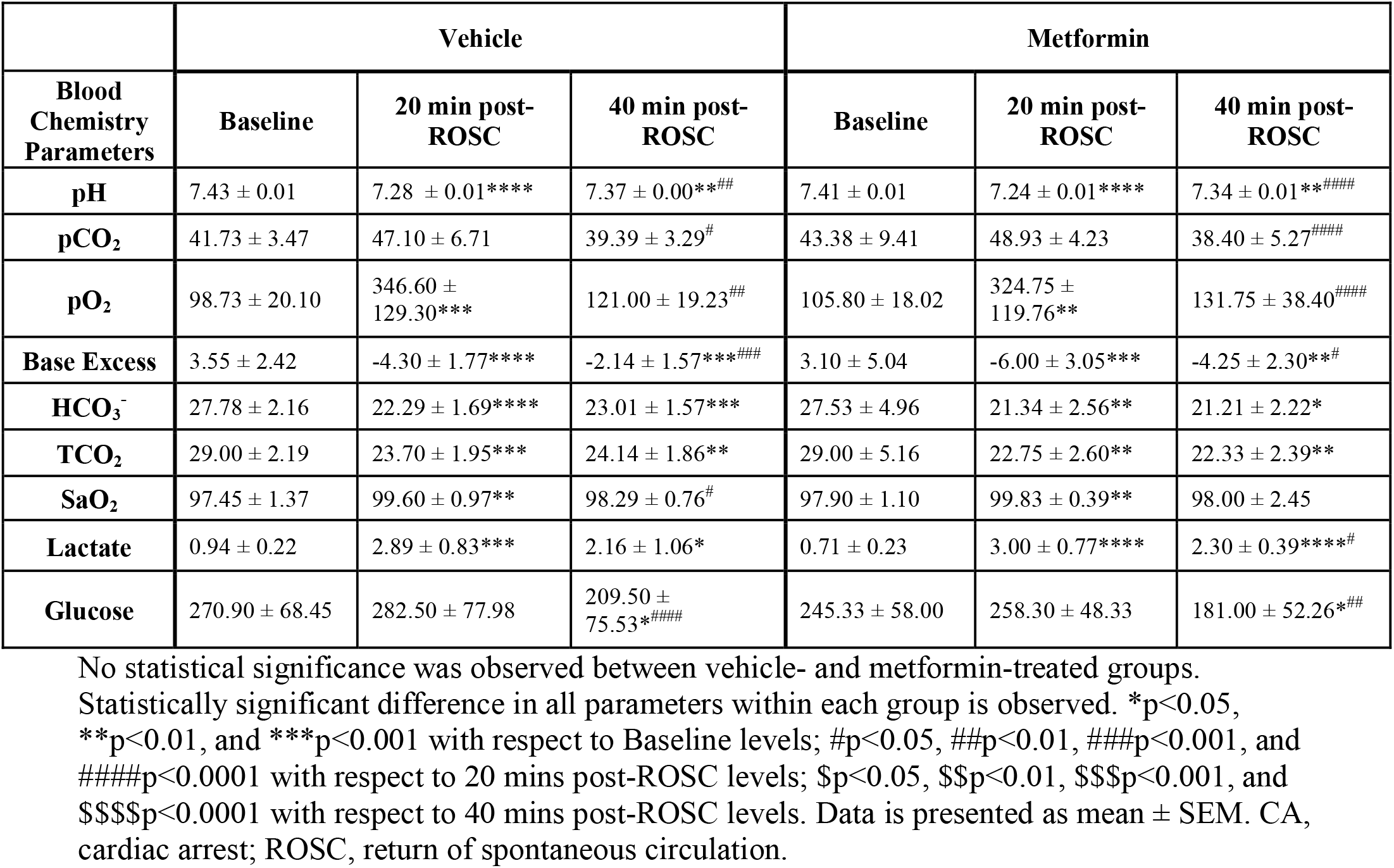
Intragroup arterial blood chemistry analysis in vehicle- and metformin-treated groups.

**Fig. 3:**
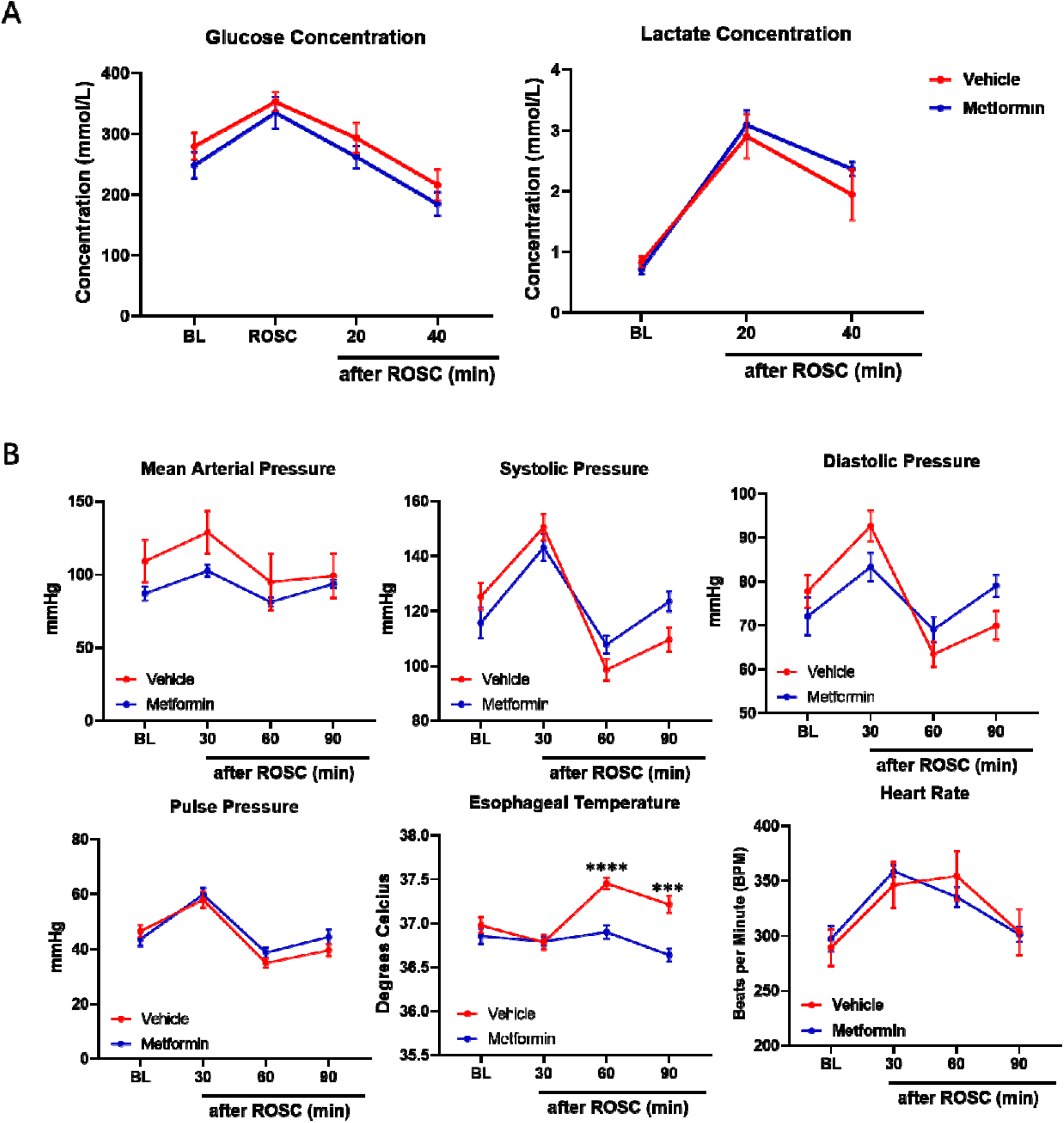
Hemodynamics, glucose, and lactate post-CA. Evaluation of plasma glucose and lactate concentration and hemodynamic parameters in vehicle- and metformin-treated groups. No statistical significance was observed between vehicle and metformin groups. ***P<0.001 and ****P<0.0001. BL, baseline; ROSC, return of spontaneous circulation.

### Metformin administration improves brain morphological features and decreases neuronal damage after 10 min CA and resuscitation

Histological analysis of the CA 1 region using Nissl and TUNEL staining for the presence of ischemic and apoptotic cells, respectively, demonstrates therapeutic benefit of metformin. A significant increase in the average number of ischemic neurons was observed in the CA 1 region in the vehicle-treated group as compared to sham animals (P<0.001), while the average number of ischemic neurons was significantly reduced after metformin treatment as compared with vehicle treatment (P<0.01; Fig. 4). Similarly, a significant increase in the average number of TUNEL-positive cells, signifying apoptosis, was observed in the hippocampus in the vehicle-treated group as compared to sham animals (P<0.001). Although still greater than sham, a significant reduction was observed in the number of apoptotic cells with metformin treatment as compared to vehicle (P<0.05). NeuN and GFAP staining was used to differentiate between neurons and astrocytes, respectively. A trend in reduced number of neurons in CA 1 region after CA was observed in the vehicle-treated group as compared with sham (P=0.08), while metformin treatment showed retention of neurons. However, there were no changes observed in astrocyte activation among the three groups. Astrocyte aggregation has been observed in the hippocampus after 7 days post-ROSC ^32^. Although there may be a time delay for astrocyte recruitment, the immediate change post-CA that leads to loss of functionality is neuronal cell death, which metformin seems to alleviate. Evaluation of the brain cellular features at 72 h post-ROSC shows that IRI is evident in the brain in terms of ischemic and apoptotic cells and that metformin treatment can mitigate the degree of damage as compared to vehicle treatment.

**Fig. 4.**
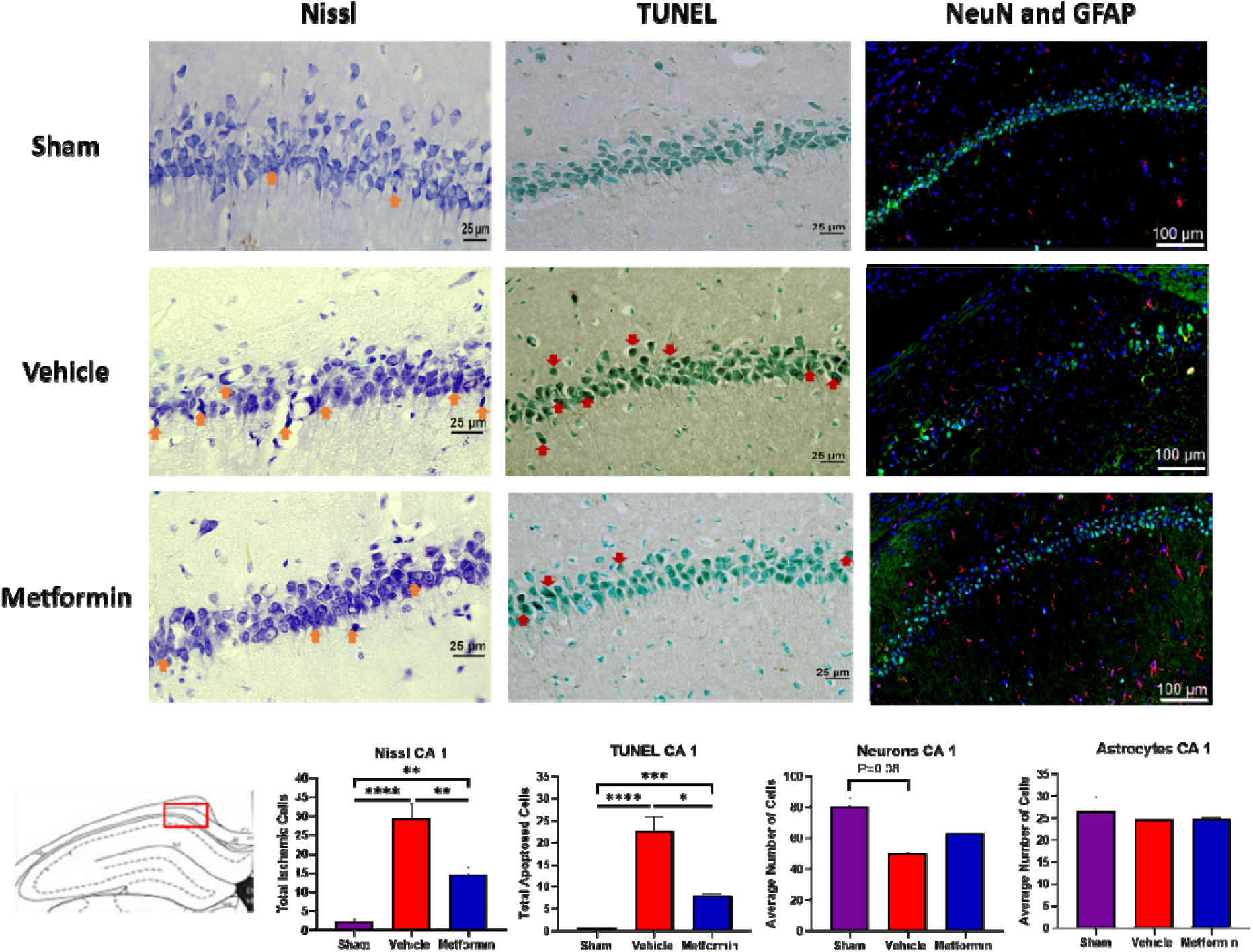
Metformin improves brain histological features post-CA. Representative Nissl, TUNEL, and NeuN/GFAP staining of hippocampus CA 1 region of sham, vehicle- and metformin-treated rats is shown with their respective cell quantitation below. Orange arrows indicate ischemic neurons, and red arrows indicate apoptotic cells. The average number of ischemic neurons and apoptotic cells in the CA1 region of the hippocampus were significantly greater in vehicle-treated rats than sham rats, and they were significantly reduced in metformin-treated rats. Merged images with NeuN/GFAP/DAPI shows a decreasing trend in NeuN-positive cells (neurons; green) in the vehicle group compared with sham, which was less severe after metformin treatment. No change was observed in GFAP-positive cells (astrocytes; red) among the three groups. DAPI (blue) indicates cellular nuclei for both neurons and astrocytes. Data is presented as mean ± SEM.*P<0.05, ** P<0.01, ***P<0.001, and ****P<0.0001.

### Metformin administration does not alter AMPK protein and mRNA levels during the early phase after resuscitation

Many studies suggest that a major target of metformin is AMPK. The reduction in oxygen and glucose due to ischemia results in decreased energy production that is sensed as a shift in the AMP/ADP:ATP ratio leading to phosphorylation of Thr172 in the α subunit increasing catabolism to help replenish the energy reserves ^33,34^. Furthermore, myristoylation and phosphorylation of the β subunit facilitates AMPK α-Thr172 activation and enzyme activity ^35,36^. As was previously shown ^11^, metformin pretreatment seems to provide protective effects against post-CA injury by modulating AMPK levels. These favorable effects also incorporate a time dimension, where chronic treatment with metformin potentiated improved outcomes. We tried to determine whether a single administration of metformin post-ROSC exerts its therapeutic benefits through AMPK or through other mechanisms. Quantification of brain protein levels of p-AMPKα, p-AMPKβ1, Total AMPKα, and Total AMPKβ1/2 and mRNA levels of AMPKα1, AMPKα2, and AMPKβ do not show any significant change among sham, vehicle- and metformin-treated rats 2h post-ROSC (Fig. 5). Our analysis of AMPK and its various subunits suggests that the role of metformin-induced AMPK activation is not observed during the early phase post-ROSC.

**Fig. 5.**
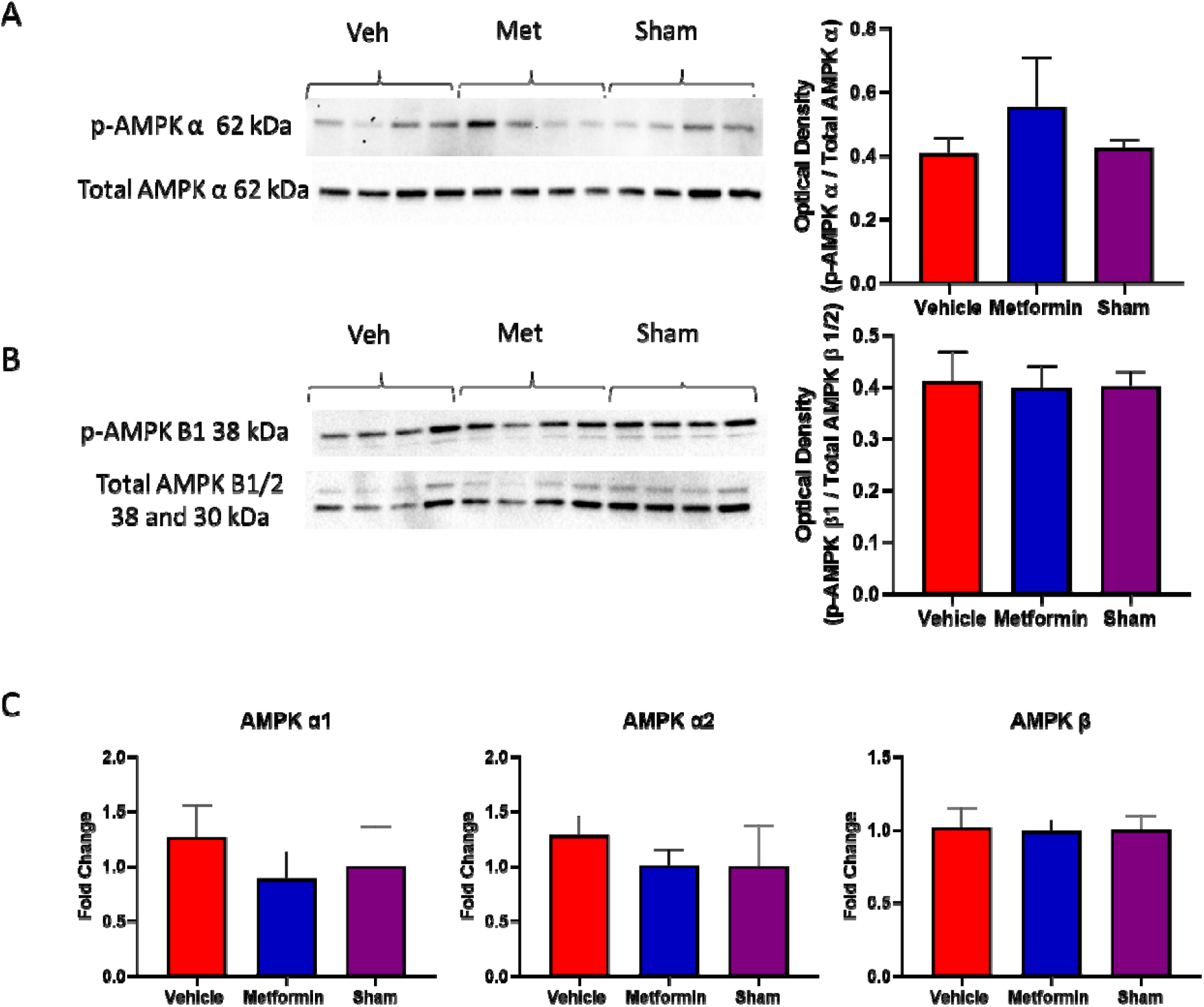
Metformin does not alter brain AMPK mRNA and protein content post-CA. Metformin treatment does not alter AMPK levels in the brain during the early phase after CA. Protein analysis of brain tissue from sham, vehicle- and metformin-treated rats 2h post-ROSC shows no difference in the levels of p-AMPKα, p-AMPKβ1, Total AMPKα, and Total AMPKβ1/2. Similarly, mRNA analysis also shows no difference in the expression of AMPKα1, AMPKα2, and AMPKβ.

### Metformin supports mitochondrial protection by preventing decrease in complex I activity, release of cytochrome c release, and oxidative stress after CA

Prior literature has suggested other, non-AMPK mechanisms for the therapeutic role of metformin ^16,37-39^. As the mitochondria is a major organelle that is severely impacted after CA, we sought to evaluate the influence of metformin on brain mitochondria. It has been well-known that metformin can directly influence complex I of the mitochondria. We analyzed the activity of complex I in whole brain homogenate and isolated brain mitochondria. Complex I activity in brain homogenate was low due to poor concentration of mitochondria; the activity did not significantly differ among sham, vehicle- and metformin-treated groups (Fig. 6). However, analysis of complex I in isolated brain mitochondria shows that vehicle-treated rats have significantly decreased complex 1 activity in isolated mitochondria when compared with sham mitochondria (P<0.05). However, metformin-treated rats do not show decreased complex I activity in isolated mitochondria suggesting metformin treatment post-CA may help to prevent decrease in complex I activity after CA. Complex IV activity was unchanged after CA. We also measured cytochrome c, a major constituent of the mitochondria that helps in ATP production. Vehicle-treated rats showed significantly decreased cytochrome c retention in isolated brain mitochondria as compared to both sham and metformin-treated rats (P<0.05) suggesting that metformin aids in protecting the mitochondria post-CA and decreasing cytochrome c release in the brain. Brain protein carbonyl concentration, which signifies oxidative damage, was significantly increased in the vehicle-treated group as compared to sham (P<0.05) and metformin treatment mitigated oxidative damage. Overall, our data suggests metformin administration post-CA is able to prevent the loss of complex I activity, maintain mitochondrial integrity as observed by retention of cytochrome c, and mitigate oxidative damage in the brain.

**Fig. 6.**
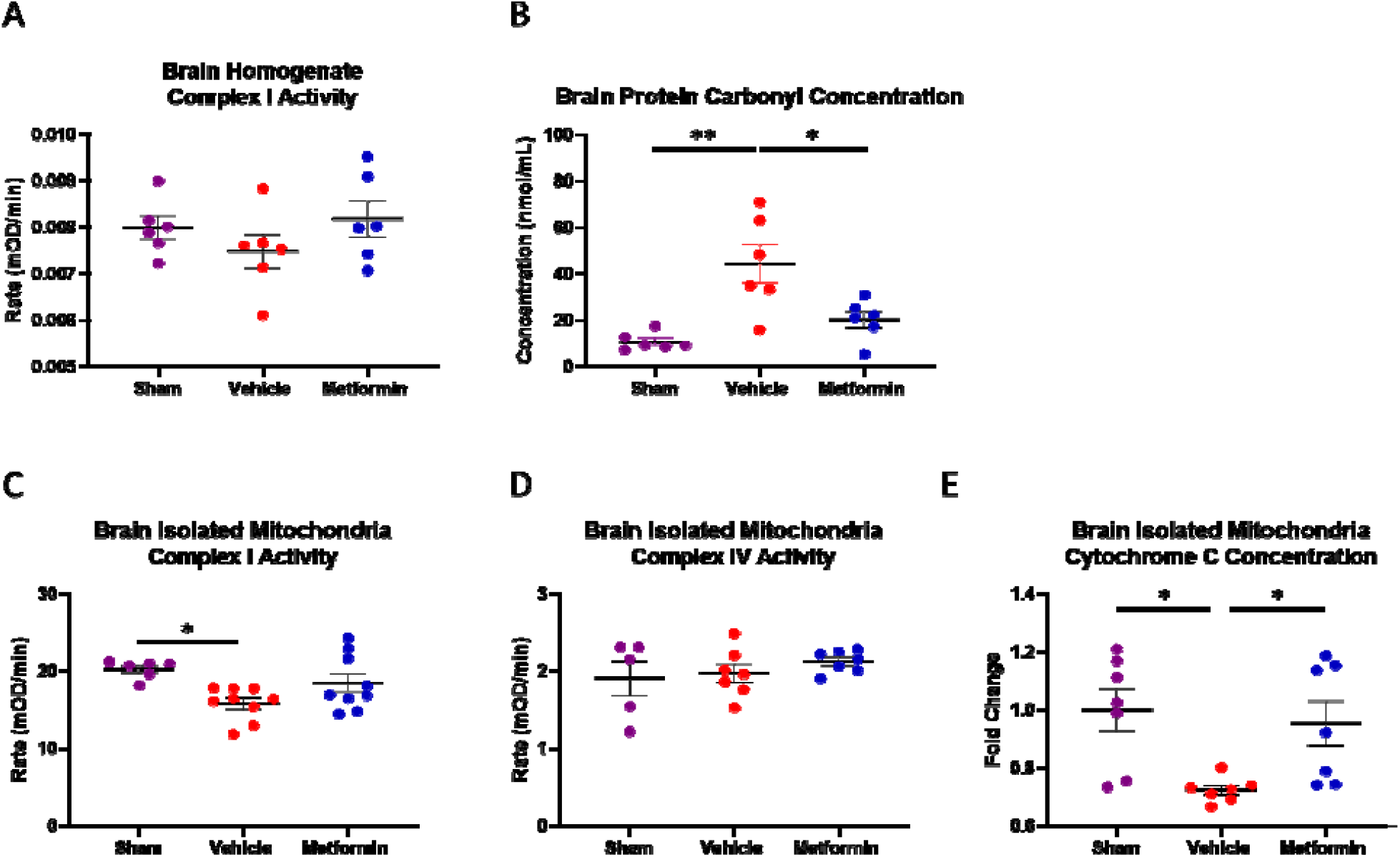
Metformin protects mitochondrial complex I activity and prevents loss of cytochrome c post-CA. Complex I activity in the total brain homogenate was low. Total brain protein carbonyl concentration, which is a surrogate marker for oxidative damage, was significantly higher in the vehicle-treated group as compared to both sham and metformin-treated group. Evaluation of complex I activity in isolated brain mitochondria demonstrated decreased activity in the vehicle-treated group as compared to sham with metformin-treated group showing normalization of activity. Complex IV activity in isolated mitochondria did not differ among the three groups. Evaluation of cytochrome c retention, a hallmark of mitochondrial and cellular health, showed severe loss of cytochrome c in the vehicle-treated group with metformin treatment preventing loss of cytochrome c from the mitochondria. Data is presented as mean ± SEM.*P<0.05; **P<0.01.

### Metformin administration potentiates earlier normalization of EEG activity after 10 min CA and resuscitation

Evaluation of brain recovery is important after CA especially when attempting to access the efficacy of a potential neuroprotective intervention. However, a critical period of neuronal damage is during the early phase after ROSC. EEG activity has already been used to assess neurological recovery in comatose patients after CA ^30^. Persistent isoelectricity, low voltage activity, or low burst-suppression patterns in EEG activity in the first 24□h predict a poor outcome, while rapid recovery towards continuous patterns within 12□h is strongly associated with better neurological outcomes ^31^. We measured EEG changes after CA up to 2 h post-ROSC to determine the therapeutic benefit of metformin during the early phase after resuscitation. Metformin treatment demonstrated an increased early bursting frequency pattern in EEG as compared with vehicle treatment (Fig. 7). The total Burst-to-Suppression Ratio (BSR), which shows the degree of neuronal firing as opposed to lack of activity, increased over time after ROSC and was significantly higher at 100 and 110 min post-ROSC after metformin treatment as compared with vehicle (P<0.05; Fig. 7C). Furthermore, the time to reach 50% of baseline BSR (95.00 ± 12.04 vs 63.33 ± 4.94 min for vehicle vs metformin; P<0.05) and the time to normalize BSR (146.20 ± 16.05 vs 92.40 ± 13.11 min for vehicle vs metformin; P<0.05) were significantly lower after metformin treatment, suggesting that metformin confers its neuroprotective effects in the early phase post-ROSC (Fig. 7, D and E).

**Fig. 7.**
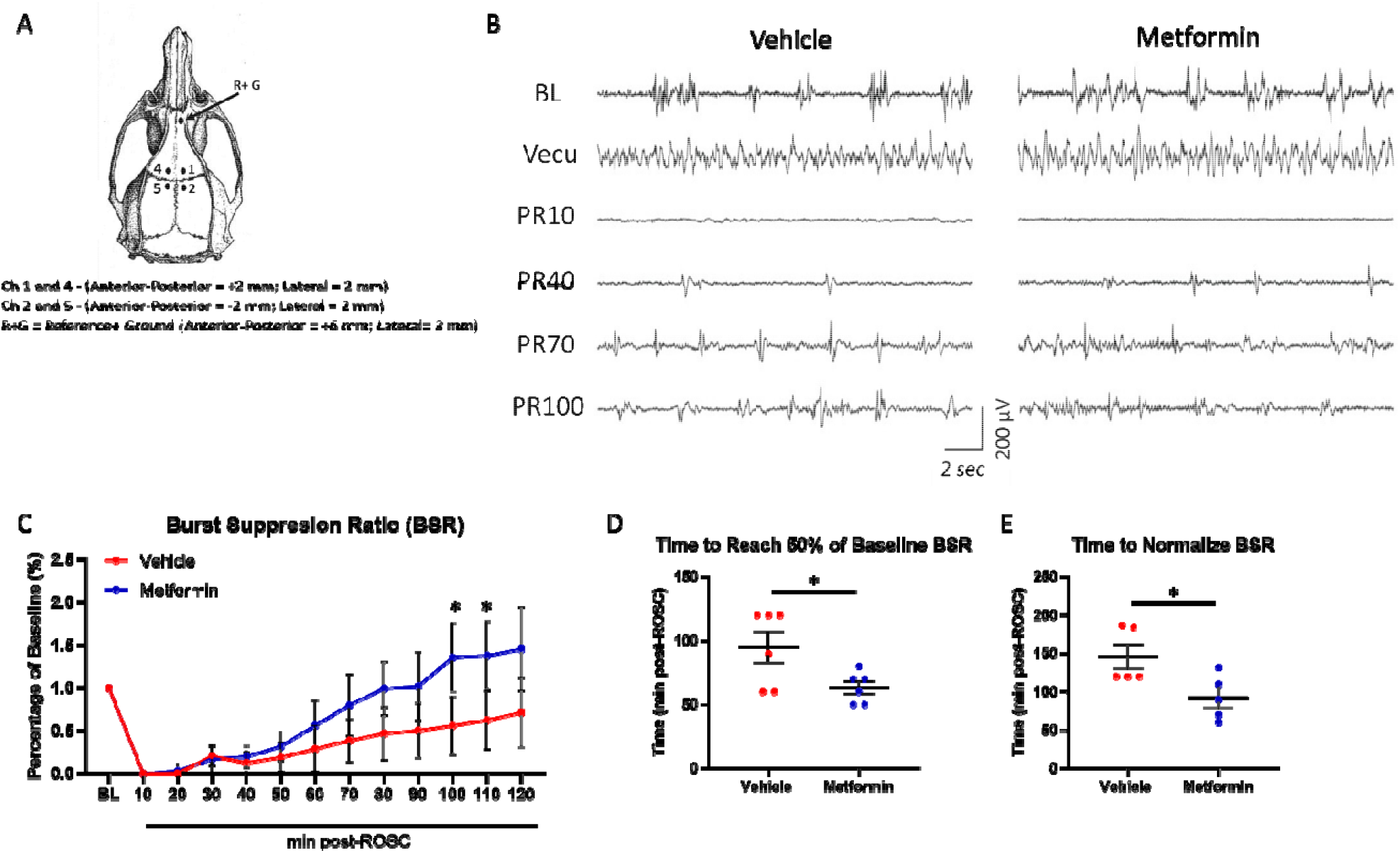
EEG post-cardiac arrest supports benefit of metformin treatment. Diagram for electrode implantation for EEG recording (A). Metformin treatment demonstrates an increased early bursting frequency pattern in EEG as compared with vehicle treatment as observed in Raw EEG (B). The total Burst-to-Suppression Ratio shows increasing trend over time after ROSC, which was significantly higher at 100 and 110 min post-ROSC after metformin treatment as compared with vehicle (C). The time to reach 50% BSR (D) and the time to normalize BSR (E) were significantly lower after metformin treatment. Data is presented as mean ± SEM.*P<0.05. BL, baseline; Vecu, vecuronium; PR10, post-ROSC 10 min; PR40, post-ROSC 40 min; PR70, post-ROSC 70 min; PR100, post-ROSC 100 min.

## Discussion

Cardiac arrest is a devastating disease process that produces whole-body IRI in which most patients do not survive ^3^. Brain damage post-CA has been implicated as a major source of poor outcomes. Furthermore, the brain mitochondria are severely injured in that their membrane lipid composition is altered ^6,40^ as well as disruption of proper function in the energy generation centers ^5,41^. As such, mitigating brain mitochondrial damage post-CA is an important avenue of exploration in developing interventions for treating CA. With is high safety profile and substantial efficacy in treating diabetes, metformin has also shown multiple beneficial effects in other diseases, such as aging, cancer, and ischemia ^12,42^. As it has been shown to interact with the mitochondria, we hypothesized that administering metformin post-CA may help mitigate mitochondrial damage in the brain and confer favorable outcomes. In our study, we found a significant increase in 72 h survival with metformin treatment as compared to vehicle (Fig. 2). Furthermore, we also observed that metformin treatment also showed significantly higher retention in neurological function as compared with vehicle treatment rats at 24 h, 48 h, and 72 h. Global IRI after CA produces continuous injury in the brain that evolves over time. It is important to evaluate the cellular damage in the brain after CA despite our positive survival findings. Therefore, our histological analysis 3 days post-CA demonstrated a substantial reduction in the number of both ischemic and apoptotic cells in the hippocampal CA 1 region of the brain when treated with metformin (Fig. 7). Our positive results suggest that metformin can be a potential therapeutic approach to treat CA.

Observing the improvement in outcomes after CA, we further wanted to explore the mechanism(s) by which these therapeutic effects are observed. However, it is noteworthy that evaluation of metformin’s mechanism is challenging to identify because of the complexity of CA and the multitude of metformin’s targets. The direct interactions of metformin and AMPK have been elusive. It is suggested that among the many mechanisms, AMPK may be activated by metformin directly or via mitochondrial pathways ^43-45^. The only attempt of using metformin in the context of CA was performed by providing metformin for 2 weeks prior to CA. It was demonstrated that metformin induces an AMPK-dependent autophagy activation to produce favorable outcomes ^11^. We hypothesized that if metformin’s therapeutic effects were in an AMPK-mediated fashion, then we should observe increased phosphorylation of both AMPK α and β even if it is provided post-CA. However, we observed that the severe IRI caused by CA did not result in major differences in the protein concentrations or mRNA levels of either the α or β subunits to activate AMPK in the brain among sham, vehicle-treated group post-CA, or metformin-treated group post-CA (Fig. 5). This suggests that although AMPK activation may be a potential mechanism for metformin’s therapeutic efficacy, this pathway may not be implicated during the early phase after ROSC.

As the role of mitochondria in CA injury becomes more established ^3,41^ and minimal changes in brain AMPK were observed, our next target was to assess the mitochondria in the brain after metformin administration post-CA. Evaluation of complex I activity in isolated brain mitochondria showed a significant decrease in complex I activity in the vehicle-treated group, which was expected and already validated in a prior study ^41^. However, metformin treatment prevented the decrease in activity (Fig. 6). Although it is presumed the metformin would inhibit mitochondrial complex I, our results suggest the opposite. The dosage and duration of treatment with metformin has a direct impact on the role it plays within the body. High local concentrations of metformin (>1mM) are considered supra-pharmacologic and can partially or fully inhibit mitochondrial complex I ^37,46^. It is noteworthy that our 100 mg/Kg dose, which would provide a maximal concentration of less than 10 mM in the whole rat, may prove to have much lower local concentrations than we expected. This would be enough to modulate rather than inhibit complex I activity. In fact, complex I shows varying effects in activity with metformin dosing, such that higher dosages can actually increase activity and have other, non-mitochondrial mechanisms of action, such as inhibition of mitochondrial glycerophosphate dehydrogenase (mGPD) ^47-50^. Therefore, the causal relationship of our significant survival outcomes post-CA and metformin’s interactions with complex I still require more exploration.

It has been shown that instead of inhibiting complex I, metformin may stabilize the deactive form of complex I that could mitigate reverse electron transport (RET)-mediated ROS generation ^16^ and direct the flow of electrons forward to facilitate mitochondrial activity. Preventing the loss of mitochondrial function, is important for maintaining cellular integrity and preventing cell death. We wanted to evaluate any evidence of early metformin protection that resulted in the reduction of apoptotic cells in the brain at 72 h (Fig. 7). We found that there was substantially decreased concentration of protein carbonylation in the brain after metformin treatment that is indicative of reduced protein oxidative damage (Fig. 6B). This result aligns with previous studies that suggest reduction in RET by metformin. Furthermore, if the proper flow of electrons is maintained, mitochondrial integrity would be preserved. We observed that there was significant improvement in the retention of cytochrome c in isolated mitochondria from rats treated with metformin post-CA (Fig. 6E). This minimize the number of cells undergoing apoptosis ^51^, resulting in decreased loss of neurons and improvement in neurological function that we observed.

Although surviving rats showed improved brain function beginning at 24 h after a single dose of metformin post-CA, we wanted to further investigate if metformin was able to confer its benefits during the early phase after ROSC. A valuable tool in investigating the influence of metformin on the brain this early is EEG. After the loss of circulation to the brain, consciousness is lost within seconds. This can be observed as a loss of EEG bursting pattern post-CA, signaling nonexistent brain activity. This pattern persists post-ROSC and its normalization can indicate a return of brain function (Fig. 7B). However, the longer the delay to normalization and/or complete lack thereof signifies severe brain damage and potential brain death ^31^. Providing metformin post-ROSC facilitates increased EEG activity and a faster time to normalization of EEG as compared with vehicle treatment (Fig. 7, C-E). This clearly demonstrates that metformin is efficacious during the early phase post-ROSC in neuroprotection. The preservation of complex I activity, reduction in brain protein oxidative damage, and the retention of cytochrome c may all lead to early normalization of brain activity. This early brain protection with metformin may be the mechanism that results in overall improved 72 h survival and brain function.

Metformin improved brain activity in the early phase post-ROSC, improved survival, and prevented neuronal cell death. Exploration of the dosage and timing of administration is a necessary next step. Optimization of the therapeutic range, especially if used in a single administration is important to prevent adverse effects that would impede survival post-CA. As many patients are on a pharmacologic regimen of daily metformin, it is important to evaluate the additive effects of an acute treatment with metformin post-CA in the pharmacologic context of chronic treatment. It is highly likely that our observed mitochondrial-mediated beneficial effects of acute treatment may still be retained post-CA when used in patients chronically treated with metformin. However, mechanistic, and therapeutic differences are important to clarify as they will better inform management for human patients. AMPK activation also may be observed with this acute on chronic treatment with metformin post-CA. Our goal was to provide metformin during the immediate phase post-ROSC, which is a critical period for sustained survival. It is possible that administering metformin either during resuscitation or at a more delayed timepoint post-ROSC may alter its benefits. Although we showed that metformin preserved complex I activity and retained cytochrome c conferring mitochondrial protection, we were unable to evaluate the mitochondrial respiratory capacity. A deeper analysis of metformin and mitochondrial interaction is required along with exploration of metformin’s relationship with other cellular organelles that have been implicated in post-CA IRI, such as the endoplasmic reticulum.

Despite some of these limitations, our study is the first to demonstrate the positive therapeutic effects of metformin administration post-CA that seem to be mediated by mitochondrial modulation rather than AMPK activation during the early phase after resuscitation in the brain. Metformin is commonly used, has a high safety profile, and seems to confer substantial benefit post-CA that can be clinically translatable. Our study supports the application of metformin as a post-CA intervention to confer survival benefits, potentiate earlier EEG activity, and protect brain function.

## Acknowledgments

We would like to thank Ms. Seyedeh Shadafarin Marashi Shoshtari, Mr. John Seunguk Baek, and Ms. Patricia Curtin for their assistance in western blotting and histological evaluation.

## Funding

This work was supported by the ZOLL Foundation, United Therapeutics, and the Emergency Department at North Shore University Hospital.

## Author contributions

M.S, R.C.C, and L.B.B conceived and designed the experiments; M.S, R.C.C., R.K.C. S.J.M, S.H., T.Y., and M.F. performed all the experiments; M.S, R.C.C., R.K.C., N.K., and S.H. wrote the original draft of the manuscript; E.P.M, S.Z., J.K, and L.B.B. supervised the study; all authors analyzed and discussed the data and have approved the final draft of the manuscript.

## Competing interests

Muhammad Shoaib, Rishabh C. Choudhary, Rupesh K. Chillale, Nancy Kim, Santiago J. Miyara, Shabirul Haque, Tai Yin, Maya Frankfurt, Ernesto P. Molmenti, Stavros Zanos, Junhwan Kim, and Lance B. Becker declare no relevant competing interests.

## Data and materials availability

All data are available in the main text or the supplementary materials.

## Notes

### Competing Interest Statement

The authors have declared no competing interest.

